# Impact of Ethylene Oxide Sterilization on PEDOT:PSS Electrophysiology Electrodes

**DOI:** 10.64898/2025.12.30.696987

**Authors:** Ali Maziz, Clement Cointe, Benjamin Reig, Christian Bergaud

## Abstract

Poly(3,4-ethylenedioxythiophene):polystyrene sulfonate (PEDOT:PSS) is widely used to fabricate conductive organic coatings for electrodes in electrophysiology. As these devices move toward clinical translation, establishing sterilization methods that preserve their functional properties is essential. Ethylene oxide (EtO) is routinely used for sterilizing heat- and moisture-sensitive medical devices due to its high penetration efficiency and low thermal load. However, the absence of systematic studies evaluating its impact on PEDOT:PSS raises concerns about the compatibility of EtO sterilization with organic electrophysiology interfaces. Here, we report the first comprehensive evaluation of EtO sterilization on PEDOT:PSS electrodes electrochemically deposited onto cortical interfaces designed for intraoperative monitoring and stimulation. EtO exposure induced only minimal changes in surface topography, with no detectable alteration of the electrical or electrochemical performance of the electrodes. Impedance spectroscopy, cyclic voltammetry, and charge-injection capacity measurements all revealed that EtO-treated electrodes retained properties comparable to untreated controls. Moreover, EtO-sterilized PEDOT:PSS coatings demonstrated robust long-term stability under accelerated lifetime testing, exhibiting negligible degradation over extended operation. These findings demonstrate that EtO sterilization is fully compatible with PEDOT:PSS-based bioelectronic interfaces and constitutes a viable pathway toward their safe and effective integration into clinical electrophysiology. This work represents an important step toward translating organic conducting polymer technologies into real-world biomedical applications.

## 1. Introduction

The development of reliable materials for long-term neural recording and stimulation remains a central challenge in neurotechnology. Conventional electrode arrays are typically fabricated using noble metals such as gold (Au), platinum (Pt), and iridium (Ir) [1], which have enabled decades of clinical applications including cochlear implants, deep brain stimulation for Parkinson’s disease, and motor function restoration [2-5]. Despite their long history of use, metal-based neural interfaces still face significant limitations. Their stiffness and mechanical mismatch with soft neural tissue contribute to chronic inflammation and scar formation, while their high intrinsic electrochemical impedance restricts safe charge injection and limits recording fidelity[6]. For chronic neural interfaces, stability of material properties and electrical performance is crucial, however, metallic materials interact poorly with surrounding tissues due to persistent mechanical, electrical, and biological mismatches [7, 8].

To address these challenges, significant effort has been devoted to developing soft, biointegrated coatings using organic and carbon-based nanomaterials. Conducting polymers (CPs) [9, 10], carbon nanotubes (CNTs) [11, 12], and graphene [13] have emerged as promising candidates for creating compliant, high-performance neural interfaces with reduced immune response and enhanced electrochemical characteristics, including low impedance, low power consumption, and high charge-storage capacity [14]. Among these, CP nanostructures, and in particular poly(3,4-ethylenedioxythiophene) doped with polystyrene sulfonate (PEDOT:PSS) [15-18], represent one of the most impactful advancements. PEDOT:PSS offers a unique combination of low impedance, high charge-injection capacity, mechanical softness, and excellent biocompatibility [19-21]. It can be easily micro- and nanopatterned [22-24], doped or functionally modified [25-28], and supports mixed ionic–electronic conduction [29, 30], enabling both efficient neural stimulation and high-quality recording of neuronal activity [31]. Consequently, PEDOT:PSS-coated electrodes exhibit superior electrical properties compared to bare metals, including higher charge-transfer capabilities at the neuroprosthetic interface and improved recording fidelity in both in vitro and in vivo studies [32-35].

As PEDOT:PSS-based devices progress toward clinical translation, ensuring robust, biocompatible sterilization becomes a fundamental requirement. Implantable medical devices must undergo complete sterilization prior to use, and common methods include autoclaving, ultraviolet (UV) irradiation, gamma irradiation, hydrogen peroxide gas plasma, and ethylene oxide (EtO). The first clinical demonstration of PEDOT:PSS-based electrocorticography (ECoG) recording, using the EtO-sterilized NeuroGrid array to capture cortical action potentials in rodents and humans has highlighted the potential of these materials for clinical neurophysiology [36]. However, the effects of sterilization on PEDOT:PSS remain poorly understood. Sterilization processes may alter the polymer’s morphology, electrical behavior, or electrochemical stability, thereby compromising device performance. To date, no systematic studies have evaluated how EtO sterilization impacts the functional properties of PEDOT:PSS electrodes, raising critical questions about the feasibility of safely translating these interfaces into clinical use.

In this work, we present the first comprehensive investigation of the influence of EtO sterilization on PEDOT:PSS-coated electrocorticography electrodes (Figure 1). We show that EtO treatment preserves the morphology of PEDOT:PSS films and induces only minimal changes in electrical and electrochemical performance. Furthermore, EtO-sterilized electrodes maintain long-term stability under accelerated lifetime testing. These findings demonstrate that EtO constitutes a compatible and reliable sterilization method for PEDOT:PSS-based neural interfaces, representing an important step toward their future adoption in clinical electrophysiology.

**Figure 1.**
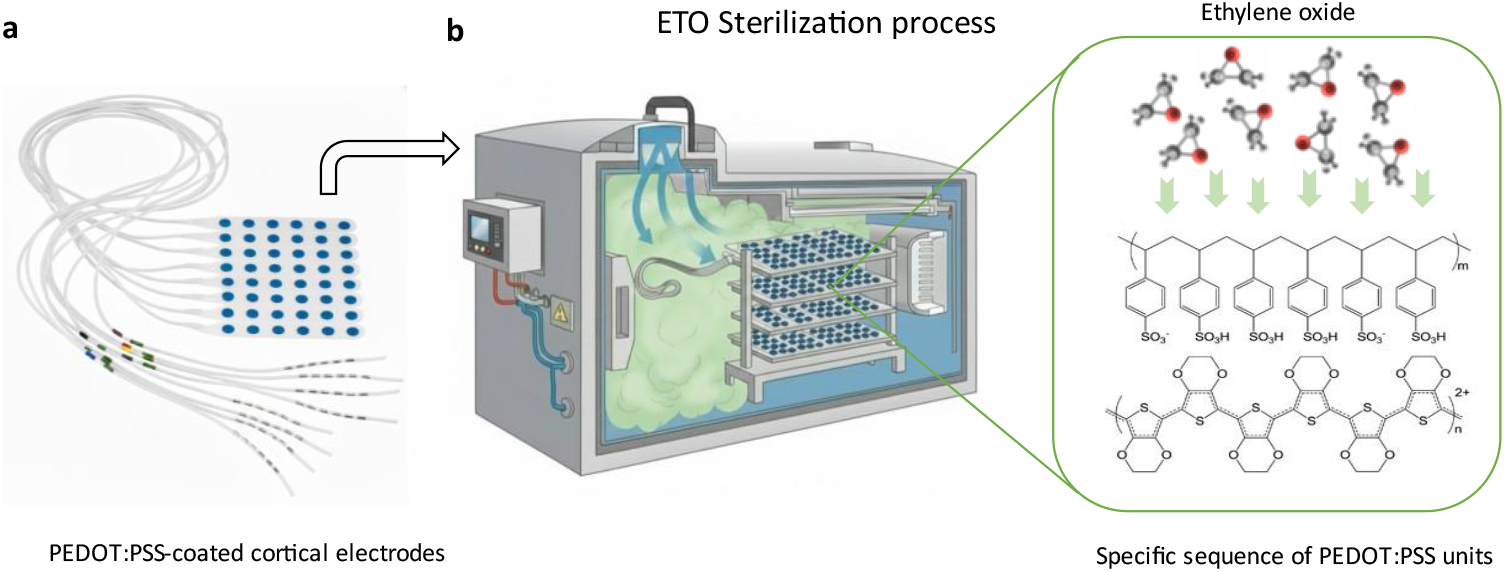
Schematic illustration of ethylene oxide (ETO) sterilization of PEDOT:PSS-coated.

## 2. Materials and Methods

### 2.1 Materials

Commercial cortical electrodes were used as reference devices in this study (DIXI Medical, cortex strip electrodes, product ID C10-06BIOM). Each flexible strip consisted of eight numbered stainless-steel contacts arranged linearly, with a contact diameter of 2.5 mm and an inter-contact spacing of 10 mm. The electrodes had a total strip width of 10 mm and a thickness below 0.8 mm, allowing close conformity to the cortical surface. The contacts were dome-shaped, and micro-perforations between strips facilitated handling and adaptation to the brain surface. The electrodes were equipped with integrated touch-proof DIN 42802 connectors, ensuring compatibility with standard clinical recording and stimulation systems.

3,4-Ethylenedioxythiophene (EDOT) and poly(sodium 4-styrenesulfonate) (NaPSS, average M_w_ ≈ 70,000) were purchased from Sigma-Aldrich and used as received. Deionized water (18 MΩ·cm) was used for the preparation of all solutions and throughout all experiments. Platinum and silver wires were purchased from World Precision Instruments (WPI).

### 2.2 Electrochemical deposition of PEDOT:PSS

The surfaces of all Pt/Ir electrodes were initially cleaned in 0.5 M H_2_SO_4_ by performing multiple electrochemical oxidation-reduction cycles. The applied potential was cycled between –0.4 V and 1.6 V versus an Ag/AgCl reference electrode, with a platinum wire serving as the counter electrode, at a scan rate of 200 mV·s^−1^ using a Biologic potentiostat (BioLogic VMP3). Following surface activation, the electrodes were rinsed with deionized water and thoroughly dried. PEDOT:PSS was subsequently deposited on the electrodes using the cyclic voltammetry (CV) technique. Each electrode underwent a single CV scan between –0.7 V and 0.9 V at a scan rate of 10 mV·s^−1^ versus Ag/AgCl. After deposition, the electrodes were rinsed with deionized water and immersed in 0.7% (w/v) NaCl aqueous for at least 30 minutes prior to further characterization.

### 2.3. Ethylene oxide sterilization of PEDOT:PSS electrodes

PEDOT:PSS cortical electrodes were sterilized using ethylene oxide (EtO) gas by an external certified provider (Steriservices). Sterilization was performed in an industrial EtO chamber with a total volume of 11 m^3^ and an effective working volume of 6.9 m^3^. The maximum mass of EtO injected per cycle was 8.0 kg, supplied as a gas mixture of 90% ethylene oxide and 10% carbon dioxide. The sterilization cycle comprised a conditioning phase of 180 minutes, followed by a gas exposure phase of 240 minutes. During exposure, the temperature was maintained between 30 and 50 °C, and relative humidity was controlled at a minimum of 50%. No post-sterilization aeration was applied. The process was designed to achieve a minimum Sterility Assurance Level (SAL) of 10^−6^.

### 2.4. Electrochemical characterizations

All electrochemical characterizations were performed using a three-electrode cell, with the microelectrodes serving as working electrodes (WEs), a thick platinum wire (2 mm diameter, ∼5 mm^2^, WPI, 99.99%) as the counter electrode (CE), and a chloritized silver wire (0.5 mm diameter) as the reference electrode (RE).

#### 2.4.1. Electrochemical Impedance Spectroscopy (EIS)

Impedance measurements were conducted by Electrochemical Impedance Spectroscopy (EIS) using a Bio-Logic VMP3 potentiostat in 0.7% (w/v) NaCl aqueous solution. Measurements were performed over the frequency range of 10 Hz to 10 kHz using a 10 mV AC signal at 0 V versus SCE.

#### 2.4.2. Cyclic voltammetry (CV)

Capacitance measurements were performed by cyclic voltammetry (CV) using the low-current channel of the Bio-Logic VMP3 potentiostat. Scans were carried out between 0,6V and -0.6 V versus Ag/AgCl in 0.7% (w/v) NaCl aqueous solution at a scan rate of 50 mV·s^−1^. Electrodes were cycled until a stable voltammogram was obtained, typically by the fourth cycle.

#### 2.4.3. Voltage transient response

To estimate the charge injection limit, voltage transient measurements were carried out at different input currents by applying charge-balanced biphasic current pulse waveforms at 10 Hz, with pulse durations of 500 µs (Figure 6d), using a Bio-Logic VSP3 potentiostat. The negative polarization potential (V_p_) was calculated by subtracting the initial access voltage (V_a_) due to solution resistance from the total voltage (V_max_). The charge injection limits were calculated by multiplying the current amplitude and pulse duration at which the polarization potential reaches the water reduction limit (−1.0 V), divided by the geometric surface area of the electrode [37, 38].

### 2.5. Scanning Electron Microscopy (SEM)

The morphology of the PEDOT:PSS coatings was examined using a HITACHI S-4800 cold field-emission high-resolution scanning electron microscope (SEM) operated at 800 V and a beam current of 2 µA.

### 2.6. Atomic Force microscopy (AFM)

AFM imaging and force spectroscopy of PEDOT:PSS deposited on Pt/Ir electrodes were performed in contact mode using a Bruker ICON system equipped with a Nanoscope V controller. SI_3_N_4_ AFM probes (MLCT, Veeco Instruments) with a pyramidal tip and an opening angle of 35° were used for all measurements. Topography and surface potential analyses of the PEDOT:PSS coatings were conducted in tapping mode and via Kelvin probe force microscopy (KFM) mapping under ambient conditions.

## 3. Results

### 3.1. PEDOT:PSS electrochemical deposition

To evaluate the impact of EtO sterilization on the electrochemical properties of PEDOT:PSS, we employed macroscopic electrocorticography (ECoG) electrodes (DIXI Medical, cortex strip electrodes, product ID C10-06BIOM). Each flexible strip consisted of eight numbered stainless-steel contacts arranged linearly, with a contact diameter of 2.5 mm and an inter-contact spacing of 10 mm, as in Figure 2b. Our study focuses on the clinical translation potential of PEDOT:PSS by examining the effects of EtO sterilization on electrode performance. This simple design allowed direct comparisons between PEDOT:PSS-modified electrodes and standard PtIr electrodes in terms of structure, morphology, electrochemical behavior, and stability, both before and after sterilization.

**Figure 2.**
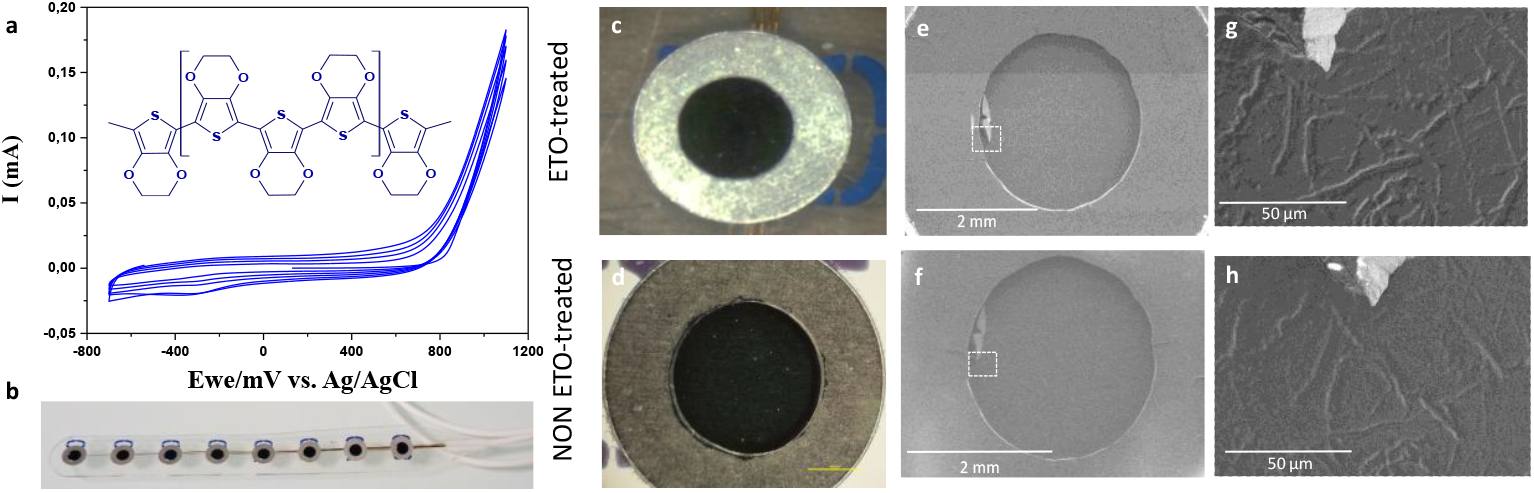
(a) Cyclic voltammograms recorded during electropolymerization on a 2.5 mm-diameter electrode (4 cycles from −0.7 to 1.1 V, scan rate: 10 mV s^−1^). (b) Optical photograph of PEDOT:PSS-coated cortical electrodes. (c,d) Optical microscopy images of PEDOT:PSS electrodes before and after ethylene oxide (ETO) sterilization, respectively. (e,f) SEM images of the same PEDOT:PSS electrode before and after ETO sterilization, respectively. (g,h) Close-up views of a localized manufacturing defect at the electrode periphery, enabling observation of the same area before (g) and after (h) ETO sterilization.

The PtIr electrodes were coated with PEDOT:PSS, a conducting polymer recognized for its excellent biocompatibility, high electrical conductivity, and superior charge-injection capabilities. We have previously demonstrated that electrochemical polymerization can deposit uniform, stable, and mechanically robust PEDOT:PSS films directly onto metal electrode sites [39]. In contrast to aqueous PEDOT:PSS dispersions typically processed into thin films, electrodeposition occurs potentiodynamically (Figure 2a), via a chain-propagation mechanism on the PtIr surface. This process produces a smooth and porous three-dimensional surface structure resulting from nucleation and growth of the polymer. The resulting PEDOT:PSS coatings had a thickness of approximately 400 nm. PEDOT:PSS electrodes subjected to EtO sterilization (Figure 1) were characterized before and after treatment to assess potential changes in electrochemical activity and surface morphology, providing a direct evaluation of sterilization compatibility for clinical applications.

### 3.2. Morphology

To assess the structural integrity of PEDOT:PSS electrodes and their interfaces with the underlying metal contacts, a critical safety consideration for clinical use, we investigated the morphological stability of the films before and after EtO sterilization. Optical microscopy revealed no visible cracking or noticeable changes in surface morphology following sterilization (Figure 2 c-d). To further examine surface stability, SEM was performed on both untreated and EtO-treated PEDOT:PSS electrodes. As shown in Figure 2 e-f, SEM images confirm that EtO sterilization does not alter the overall morphology of the polymer surface, consistent with optical observations. A minor manufacturing defect was present at the periphery of one electrode, allowing for direct comparison of the same area before (Figure 2g) and after (Figure 2h) sterilization. Importantly, this defect did not propagate to other electrodes. The surface of EtO-treated PEDOT:PSS appeared relatively smooth, while the untreated films displayed slightly less uniform structures.

AFM further quantified the surface topography. As shown in Figure 3, EtO-treated PEDOT:PSS exhibited a slightly smoother surface compared to untreated films. Nonetheless, the overall surface roughness was not significantly different, with a root mean square roughness of 71 nm for untreated PEDOT:PSS and 68 nm for EtO-treated films. These results indicate that EtO sterilization preserves the morphological integrity of PEDOT:PSS coatings, supporting their suitability for clinical applications.

**Figure 3.**
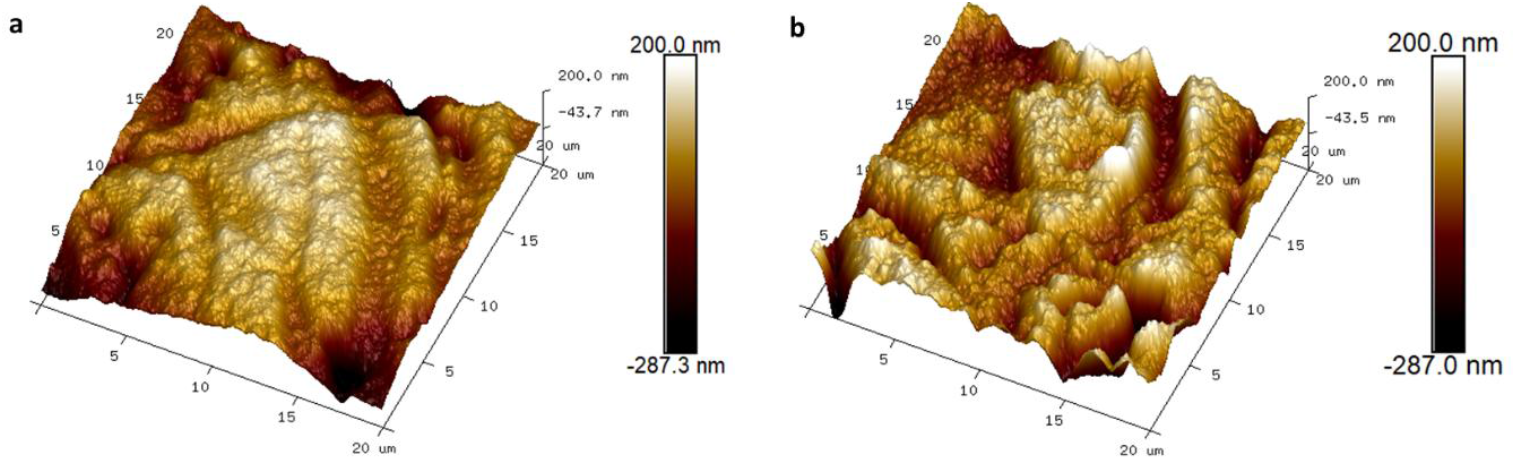
AFM topography images of PEDOT:PSS over a 20 × 20 μm^2^ area before (a) and after (b) ethylene oxide sterilization.

### 3.2. Electrochemical Impedance Spectroscopy

We next evaluated the effect of EtO sterilization on the electrical and electrochemical properties of PEDOT:PSS electrodes using electrochemical impedance spectroscopy (EIS), cyclic voltammetry (CV), and charge storage capacity measurements. Assessing potential alterations in electrode performance after ETO sterilization is critical to ensure the functional reliability of PEDOT:PSS coatings. Non-coated PtIr electrodes of comparable diameter were used as controls.

Figure 4 show Bode plots of impedance magnitude and phase angle for electrodes before and after sterilization, for both PEDOT:PSS-coated and bare PtIr electrodes. Across the physiologically relevant frequency range (10 Hz–10 kHz), PEDOT:PSS electrodes exhibited substantially lower impedance than PtIr electrodes, consistent with their enhanced electrochemical properties. After EtO sterilization, PEDOT:PSS electrodes showed only minor increases in impedance, with the average value at 100 Hz rising from 195 ± 4 Ω to 211 ± 7 Ω (Figure 4c). Bare PtIr electrodes displayed a similar trend, with average impedance increasing from 1270 ± 340 Ω to 1810 ± 188 Ω. The phase angle also increased across all frequencies following sterilization, consistent with the observed impedance changes (Figure 4b). To further quantify these effects, we calculated the relative change in impedance at 10 Hz, 100 Hz, and 1 kHz, as well as across the full 10 Hz–10 kHz spectrum, presented as Z_post_/Z_pre_. These analyses reveal that the largest impact of EtO sterilization occurs at low frequencies, where the ratio Z_post_/Z_pre_ was 1.31 ± 0.08 at 10 Hz. At 100 Hz and 1 kHz, the corresponding ratios were 1.08 ± 0.03 and 1.04 ± 0.03, respectively. Overall, these results indicate that EtO sterilization induces only minor changes in PEDOT:PSS impedance, particularly at frequencies relevant for neural recording and stimulation, supporting the compatibility of this sterilization method for clinical applications.

**Figure 4.**
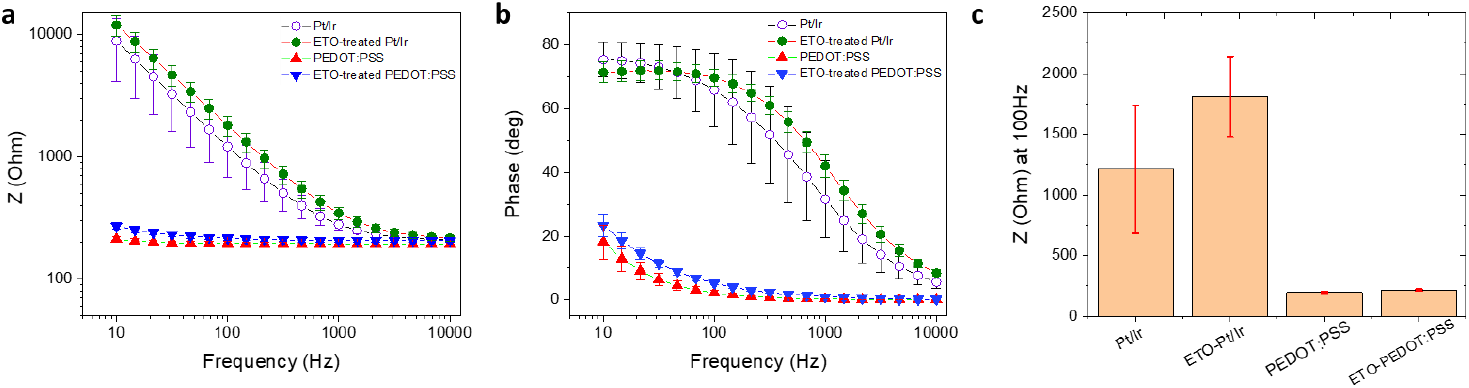
Electrochemical comparison of PEDOT:PSS, ETO-treated PEDOT:PSS, PtIr, and ETO-treated PtIr electrodes (D = 2.5 mm). (a,b) Impedance magnitude and phase angle, respectively, measured in 0.7% (w/v) NaCl aqueous solution. (c) Impedance at 100 Hz for PEDOT:PSS, ETO-treated PEDOT:PSS, PtIr, and ETO-treated PtIr electrodes.

### 3.3 Charge Storage capacity (CSC)

We next investigated the effect of EtO sterilization on the charge storage capacity (CSC) of PEDOT:PSS electrodes, a key metric of their ability to deliver charge during neural stimulation. Cyclic voltammetry (CV) was performed in 0.7% (w/v) NaCl aqueous within a potential window of −0.6 to +0.6 V (Figure 5a). The cathodal (CSCc) and anodal (CSCa) charge storage capacities were calculated by integrating the cathodal and anodal currents, respectively (Figure 5b). Histograms of CSC values (Figure 5c-d) show minor differences between EtO-treated and untreated PEDOT:PSS electrodes. EtO sterilization resulted in a modest decrease of approximately 9% in CSCc and 7% in CSCa. This slight reduction reflects a small decrease in the current response to voltage ramping, but the overall charge storage ability of the electrodes remained substantially higher than that of bare PtIr controls. Specifically, PtIr electrodes exhibited average CSC values of 3.2 mC·cm^−2^ (CSCc) and 0.98 mC·cm^−2^ (CSCa), whereas PEDOT:PSS-coated electrodes achieved 10.2 mC·cm^−2^ (CSCc) and 9.2 mC·cm^−2^ (CSCa).

**Figure 5.**
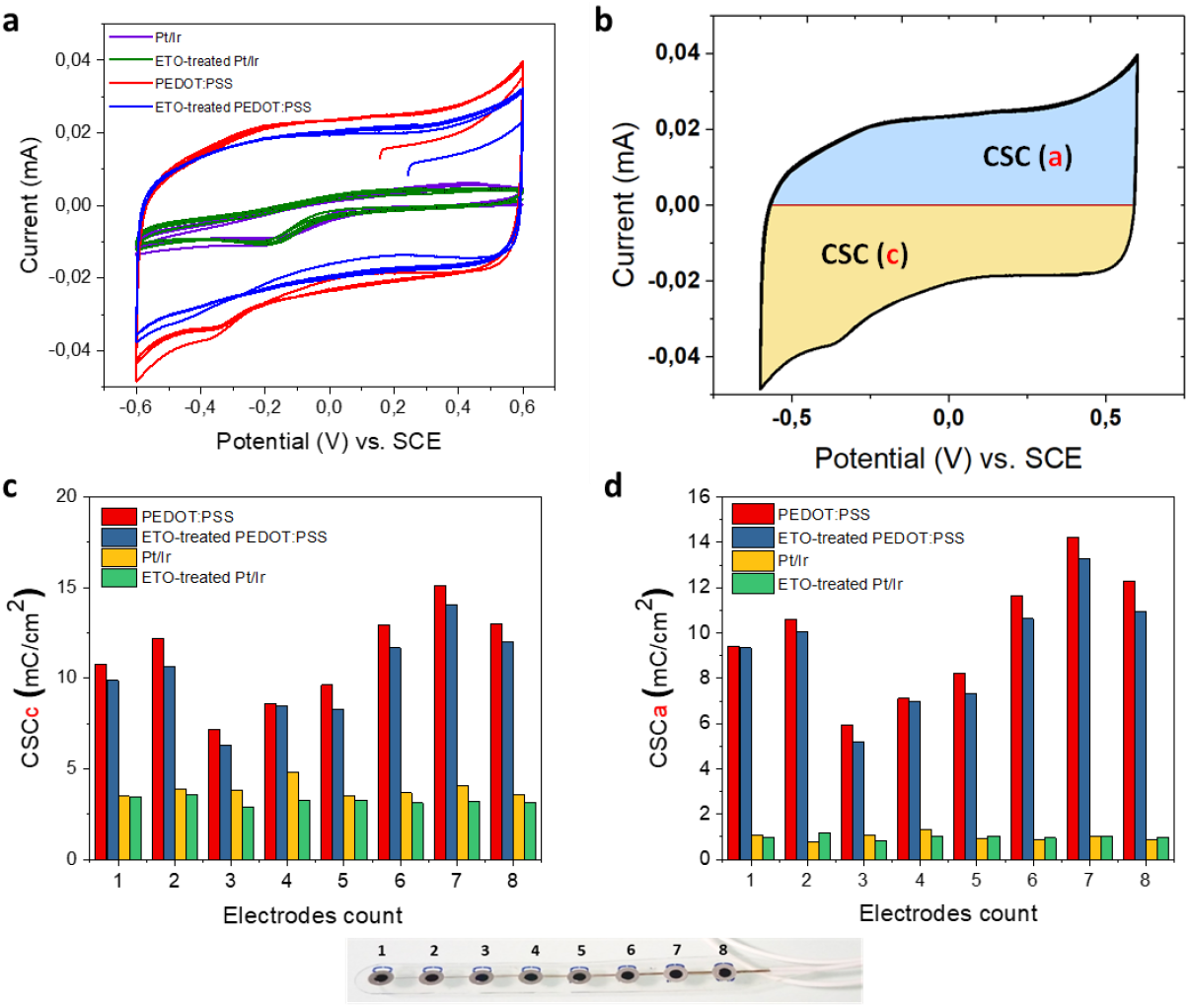
(a) Cyclic voltammograms recorded during electrochemical characterization on a 2.5 mm-diameter electrode (4 cycles from −0.6 to 0.6 V, scan rate: 200 mV·s^−1^) in 0.7% (w/v) NaCI aqueous solution, (b) Cathodal (CSCc) and anodal (CSCa) charge storage capacities, calculated by integrating the cathodal and anodal currents, respectively. (c,d) Histograms of CSCc and CSCa values, respectively, obtained from eight electrodes on the same strip.

The marked improvement in charge storage is attributed to the high surface area and porous structure of the PEDOT:PSS coating, which facilitates efficient electrolyte ion diffusion. Importantly, despite the minor reduction after EtO sterilization, PEDOT:PSS electrodes maintain a substantial electrochemical advantage over metallic electrodes, demonstrating their robustness and suitability for clinical neural interface applications.

### 3.4. Electrical stimulation

While CSC provides a measure of the total charge an electrode can store under slow voltage ramps, it does not fully represent performance under neural stimulation conditions, which typically involve cathodal-first, biphasic current pulses of millisecond duration. During such fast stimulation, only a fraction of the total CSC is accessible. Therefore, we assessed the charge injection limit (CIL) of PEDOT:PSS electrodes to evaluate their functional performance before and after EtO sterilization. CIL is defined as the maximum charge that can be injected into the solution during a stimulation pulse without exceeding the water electrolysis limits. To determine safe polarization levels, the water window of PEDOT:PSS electrodes was measured in 0.7% (w/v) NaCl aqueous solution using cyclic voltammetry at 200 mV/s versus an Ag/AgCl reference electrode (Figure 6a). Water reduction and oxidation potentials were found at −1 V and 0.6 V, respectively. Based on these limits, a conservative polarization potential of −0.85 V was selected as the safe cathodic polarization threshold, ensuring operation within the water window and avoiding the onset of water electrolysis. Next, the voltage excursions in response to biphasic, cathodic first, current pulses were recorded with a 500 µs pulse width in 0.7% (w/v) NaCl (Figure 6b-c). Using a range of pulse current intensities, we defined the CIL as the amount of charge injected which caused polarization of the electrode beyond its water hydrolysis window (Figure 6d). Figure 6e compares the electrode V_p_ measured during cathodal-first biphasic current pulses for untreated and EtO-treated PEDOT:PSS microelectrodes. For a given injected current, EtO-treated PEDOT:PSS electrodes exhibit a slightly more negative polarization potential, resulting in a modestly higher absolute V_p_ compared to untreated electrodes. This indicates a marginally increased voltage excursion during fast current pulsing following EtO sterilization. Importantly, despite this small increase in polarization, the V_p_ values for EtO-treated electrodes remain well within the defined water window, with the safe polarization limit set at -0.85 V just prior to water reduction. Consequently, the observed shift does not compromise electrochemical safety. Instead, it translates into only a minor reduction in the calculated charge injection, as shown in Figure 6f. The calculated CIL for PEDOT:PSS and ETO-treated PEDOT:PSS electrodes was 0.59 ± 0.2 mC/cm^2^ and 0.50 ± 0.1 mC/cm^2^ respectively. Overall, the results demonstrate that EtO sterilization induces only minimal changes in the interfacial charge-transfer dynamics of PEDOT:PSS under sub-millisecond stimulation conditions, preserving the electrode’s ability to inject charge efficiently and safely.

**Figure 6.**
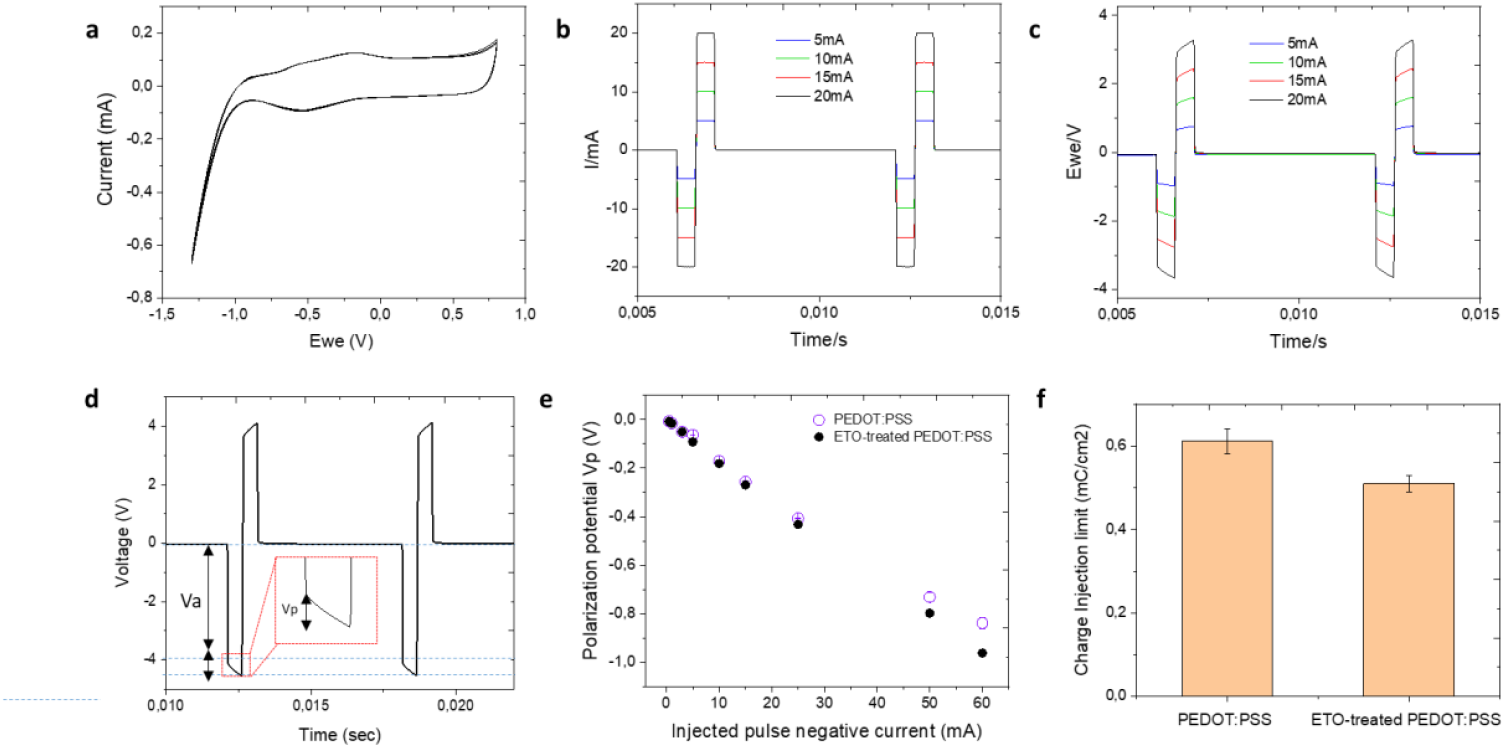
*In vitro* biphasic stimulation assessment. (a) Determination of the water reduction potential by cyclic voltammetry (CV) of a PEDOT:PSS electrode composite in 0.7% (w/v) NaCl aqueous solution at a scan rate of 200 mV·s^−1^ versus Ag/AgC . (b) Biphasic charge-balanced current pulses and (c) corresponding voltage responses recorded at different charge injection levels ranging from 5 to 20 mA. (d) Determination of the negative polarization potential (V_p_) by subtracting the initial access voltage (V_a_), arising from solution resistance, from the maximum voltage (V_max_). (e) Polarization potentials (V_p_) measured at different current pulse amplitudes. (f) Comparison of the charge injection limit (CIL) values between PEDOT:PSS and ETO-treated PEDOT:PSS electrodes.

### 3.5 Long-Term Stability of EtO-Treated PEDOT:PSS Electrodes

To evaluate the long-term stability of EtO-treated PEDOT:PSS electrodes, the devices were immersed in 0.7% (w/v) NaCl aqueous solution at 37°C for 34 days. EIS measurements (10 Hz–10 kHz) were performed approximately every two days. Figure 7 presents the impedance evolution at 100 Hz for all electrodes over the course of the study. The initial impedance of PEDOT:PSS electrodes measured at room temperature (20°C) in ACSF was 177 ± 10 Ω. Upon incubation at 37°C, the median impedance decreased to 161 ± 5 Ω, consistent with enhanced mobility of charge carriers at physiological temperature. Throughout the 34-day period, the electrodes exhibited remarkable stability, with median impedance values at 100 Hz remaining around 159 ± 5 Ω at 37°C. These results demonstrate that EtO-treated PEDOT:PSS electrodes maintain stable electrochemical properties under prolonged exposure to physiological conditions, highlighting their robustness and suitability for long-term neural interface applications.

**Figure 7.**
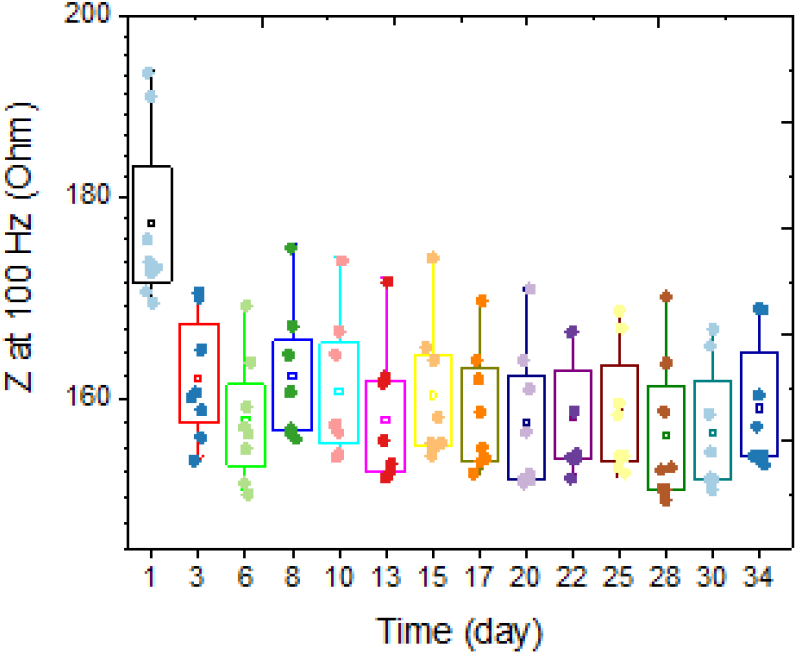
Evaluation of the stability of ETO-treated PEDOT:PSS electrodes. Electrodes were immersed in in 0.7% (w/v) NaCl aqueous solution at 37°C for 34 days. Electrode impedance was monitored using a platinum wire as the counter electrode and an Ag/AgCl as the reference electrode. Box-and-whisker plots show the evolution of impedance at 100 Hz for eight PEDOT:PSS electrodes as a function of soaking time.

## 4. Discussion

The clinical translation of PEDOT:PSS-based neural interfaces requires sterilization methods that preserve the structural, electrical, and electrochemical properties of the polymer coatings. EtO sterilization is widely used for heat- and moisture-sensitive medical devices, yet its compatibility with CPs has remained largely unexplored. In this work, we provide the first evaluation of the effects of EtO sterilization on electrochemically deposited PEDOT:PSS electrodes intended for clinical electrophysiology, demonstrating that EtO treatment preserves both morphological integrity and functional performance.

Morphological analyses reveal that EtO sterilization does not induce macroscopic or microscopic damage to PEDOT:PSS coatings (Figure 2). Optical microscopy and SEM imaging show no evidence of cracking, delamination, or film discontinuities after sterilization. AFM measurements further confirm that surface roughness remains essentially unchanged, with only a slight smoothing of the PEDOT surface observed after EtO exposure (Figure 3). This minor change may reflect subtle polymer chain relaxation or limited surface reorganization induced during the sterilization process, potentially facilitated by transient exposure to reactive gases or residual moisture. Importantly, these effects do not compromise film integrity or adhesion to the underlying PtIr substrate, which is critical for ensuring safety and reliability in clinical electrodes.

EIS demonstrates that EtO sterilization has only a minimal impact on the electrical properties of PEDOT:PSS electrodes. A modest increase in impedance is observed, particularly at low frequencies, with an average increase of approximately 8% at 100 Hz (Figure 4). This frequency range is most sensitive to interfacial phenomena, suggesting that EtO sterilization may slightly alter ion transport or interfacial capacitance at the polymer-electrolyte interface. Nevertheless, PEDOT:PSS electrodes retain impedance values that are substantially lower than those of bare PtIr electrodes across the entire frequency spectrum relevant for neural recording and stimulation. From a practical perspective, the post-sterilization impedance remains well within the optimal range for high-quality electrophysiological measurements. Consistent with the impedance results, cyclic voltammetry reveals only a modest reduction in charge storage capacity following EtO sterilization. The observed decreases of approximately 9% in cathodal CSC and 7% in anodal CSC remain small compared to the large enhancement provided by PEDOT:PSS relative to metallic electrodes (Figure 5). These changes may arise from slight reductions in electroactive surface area or minor alterations in redox accessibility within the polymer matrix. However, the overall electrochemical reversibility and capacitive behavior of the PEDOT:PSS films are preserved, indicating that EtO sterilization does not induce significant chemical degradation or loss of electroactivity. Importantly, functional stimulation performance, assessed through charge injection limit measurements under physiologically relevant pulsed conditions, is largely unaffected by EtO sterilization (Figure 6). The small (∼14%) decrease in CIL remains within experimental variability and does not meaningfully restrict the safe stimulation window. Because CIL directly determines the maximum charge that can be delivered without triggering irreversible electrochemical reactions such as water electrolysis, preservation of high CIL values is essential for clinical neural stimulation. These results confirm that EtO-sterilized PEDOT:PSS electrodes maintain their ability to safely deliver sub millisecond-scale biphasic current pulses. Long-term stability under physiological conditions is another critical requirement for clinical neural interfaces. Accelerated aging experiments at 37°C show that EtO-treated PEDOT:PSS electrodes exhibit highly stable impedance over more than one month of continuous immersion (Figure 7). This stability indicates that EtO sterilization does not introduce latent defects or chemical instabilities that could compromise long-term performance in vivo.

Taken together, these findings demonstrate that EtO sterilization is fully compatible with electrochemically deposited PEDOT:PSS electrodes. Unlike sterilization methods such as gamma irradiation or high-temperature autoclaving, which can induce polymer chain scission, oxidation, or delamination, EtO preserves the mixed ionic–electronic conduction properties that underlie the superior electrochemical performance of PEDOT:PSS. This study fills a critical gap in the literature by providing quantitative, multi-modal evidence supporting the use of EtO sterilization for organic conducting polymer-based neural interfaces.

## 5. Conclusion

In this study, we provide a comprehensive evaluation of the effects of EtO sterilization on PEDOT:PSS-coated electrophysiology electrodes. We demonstrate that electrochemically deposited PEDOT:PSS on PtIr electrodes can be effectively sterilized using EtO without appreciable alterations to film morphology. Electrical and electrochemical properties, including impedance, charge storage capacity, and charge injection limit, were only minimally affected, and long-term stability in physiological conditions was preserved. These findings establish that EtO sterilization, a method widely available in clinical laboratories is fully compatible with PEDOT:PSS-based biomedical devices. By confirming the structural, electrochemical, and functional integrity of PEDOT:PSS electrodes after sterilization, this work represents a crucial step toward the safe translation of organic conducting polymer interfaces into clinical neuroengineering applications. While this work focuses on macroscopic ECoG electrodes, the conclusions are expected to be broadly applicable to PEDOT:PSS-coated microelectrode arrays and other bioelectronic devices fabricated by electrochemical deposition. Future studies may explore the effects of repeated sterilization cycles, alternative PEDOT formulations, or molecular-level chemical changes induced by EtO exposure. Nonetheless, the present results establish a robust and clinically relevant foundation for the safe translation of PEDOT:PSS-based electrophysiology interfaces.

## References

1. Cogan, S. F., Neural stimulation and recording electrodes. Annu. Rev. Biomed. Eng. 2008, 10, (1), 275–309.

2. Chapin, J. K.; Moxon, K. A.; Markowitz, R. S.; Nicolelis, M. A., Real-time control of a robot arm using simultaneously recorded neurons in the motor cortex. Nature neuroscience 1999, 2, (7), 664–670.

3. Wessberg, J.; Stambaugh, C. R.; Kralik, J. D.; Beck, P. D.; Laubach, M.; Chapin, J. K.; Kim, J.; Biggs, S. J.; Srinivasan, M. A.; Nicolelis, M. A., Real-time prediction of hand trajectory by ensembles of cortical neurons in primates. Nature 2000, 408, (6810), 361–365.

4. Hatsopoulos, N. G.; Donoghue, J. P., The science of neural interface systems. Annual review of neuroscience 2009, 32, (1), 249–266.

5. Benabid, A. L.; Chabardes, S.; Mitrofanis, J.; Pollak, P., Deep brain stimulation of the subthalamic nucleus for the treatment of Parkinson’s disease. The Lancet Neurology 2009, 8, (1), 67–81.

6. Sharafkhani, N.; Kouzani, A. Z.; Adams, S. D.; Long, J. M.; Lissorgues, G.; Rousseau, L.; Orwa, J. O., Neural tissue-microelectrode interaction: Brain micromotion, electrical impedance, and flexible microelectrode insertion. Journal of Neuroscience Methods 2022, 365, 109388.

7. Polikov, V. S.; Tresco, P. A.; Reichert, W. M., Response of brain tissue to chronically implanted neural electrodes. Journal of neuroscience methods 2005, 148, (1), 1–18.

8. Salatino, J. W.; Ludwig, K. A.; Kozai, T. D.; Purcell, E. K., Glial responses to implanted electrodes in the brain. Nature biomedical engineering 2017, 1, (11), 862–877.

9. Green, R. A.; Lovell, N. H.; Wallace, G. G.; Poole-Warren, L. A., Conducting polymers for neural interfaces: challenges in developing an effective long-term implant. Biomaterials 2008, 29, (24-25), 3393–3399.

10. Asplund, M.; Nyberg, T.; Inganäs, O., Electroactive polymers for neural interfaces. Polymer Chemistry 2010, 1, (9), 1374–1391.

11. Keefer, E. W.; Botterman, B. R.; Romero, M. I.; Rossi, A. F.; Gross, G. W., Carbon nanotube coating improves neuronal recordings. Nature nanotechnology 2008, 3, (7), 434–439.

12. Lovat, V.; Pantarotto, D.; Lagostena, L.; Cacciari, B.; Grandolfo, M.; Righi, M.; Spalluto, G.; Prato, M.; Ballerini, L., Carbon nanotube substrates boost neuronal electrical signaling. Nano letters 2005, 5, (6), 1107–1110.

13. Lu, Y.; Liu, X.; Kuzum, D., Graphene-based neurotechnologies for advanced neural interfaces. Current Opinion in Biomedical Engineering 2018, 6, 138–147.

14. Kotov, N. A.; Winter, J. O.; Clements, I. P.; Jan, E.; Timko, B. P.; Campidelli, S.; Pathak, S.; Mazzatenta, A.; Lieber, C. M.; Prato, M., Nanomaterials for neural interfaces. Advanced Materials 2009, 21, (40), 3970–4004.

15. Cui, X.; Martin, D. C., Electrochemical deposition and characterization of poly (3, 4-ethylenedioxythiophene) on neural microelectrode arrays. Sensors and Actuators B: Chemical 2003, 89, (1-2), 92–102.

16. Yang, J.; Kim, D. H.; Hendricks, J. L.; Leach, M.; Northey, R.; Martin, D. C., Ordered surfactant-templated poly (3, 4-ethylenedioxythiophene)(PEDOT) conducting polymer on microfabricated neural probes. Acta Biomaterialia 2005, 1, (1), 125–136.

17. Ludwig, K. A.; Uram, J. D.; Yang, J.; Martin, D. C.; Kipke, D. R., Chronic neural recordings using silicon microelectrode arrays electrochemically deposited with a poly (3, 4-ethylenedioxythiophene)(PEDOT) film. Journal of neural engineering 2006, 3, (1), 59.

18. Ludwig, K. A.; Langhals, N. B.; Joseph, M. D.; Richardson-Burns, S. M.; Hendricks, J. L.; Kipke, D. R., Poly (3, 4-ethylenedioxythiophene)(PEDOT) polymer coatings facilitate smaller neural recording electrodes. Journal of neural engineering 2011, 8, (1), 014001.

19. Abidian, M. R.; Martin, D. C., Multifunctional nanobiomaterials for neural interfaces. Advanced functional materials 2009, 19, (4), 573–585.

20. Poole-Warren, L.; Lovell, N.; Baek, S.; Green, R., Development of bioactive conducting polymers for neural interfaces. Expert review of medical devices 2010, 7, (1), 35–49.

21. Yoon, H.; Jang, J., Conducting-polymer nanomaterials for high-performance sensor applications: issues and challenges. Advanced Functional Materials 2009, 19, (10), 1567–1576.

22. Gomez, N.; Lee, J. Y.; Nickels, J. D.; Schmidt, C. E., Micropatterned polypyrrole: a combination of electrical and topographical characteristics for the stimulation of cells. Advanced functional materials 2007, 17, (10), 1645–1653.

23. Khaldi, A.; Falk, D.; Bengtsson, K.; Maziz, A.; Filippini, D.; Robinson, N. D.; Jager, E. W., Patterning highly conducting conjugated polymer electrodes for soft and flexible microelectrochemical devices. ACS Applied Materials & Interfaces 2018, 10, (17), 14978–14985.

24. Vajrala, V. S.; Elkhoury, K.; Pautot, S.; Bergaud, C.; Maziz, A., Hollow ring-like flexible electrode architecture enabling subcellular multi-directional neural interfacing. Biosensors and Bioelectronics 2023, 227, 115182.

25. Green, R. A.; Lovell, N. H.; Poole-Warren, L. A., Cell attachment functionality of bioactive conducting polymers for neural interfaces. Biomaterials 2009, 30, (22), 3637–3644.

26. Green, R. A.; Hassarati, R. T.; Bouchinet, L.; Lee, C. S.; Cheong, G. L.; Yu, J. F.; Dodds, C. W.; Suaning, G. J.; Poole-Warren, L. A.; Lovell, N. H., Substrate dependent stability of conducting polymer coatings on medical electrodes. Biomaterials 2012, 33, (25), 5875–5886.

27. King, Z. A.; Shaw, C. M.; Spanninga, S. A.; Martin, D. C., Structural, chemical and electrochemical characterization of poly (3, 4-Ethylenedioxythiophene)(PEDOT) prepared with various counter-ions and heat treatments. Polymer 2011, 52, (5), 1302–1308.

28. Abidian, M. R.; Daneshvar, E. D.; Egeland, B. M.; Kipke, D. R.; Cederna, P. S.; Urbanchek, M. G., Hybrid conducting polymer–hydrogel conduits for axonal growth and neural tissue engineering. 2012.

29. Rivnay, J.; Inal, S.; Collins, B. A.; Sessolo, M.; Stavrinidou, E.; Strakosas, X.; Tassone, C.; Delongchamp, D. M.; Malliaras, G. G., Structural control of mixed ionic and electronic transport in conducting polymers. Nature communications 2016, 7, (1), 11287.

30. Maziz, A.; Özgür, E.; Bergaud, C.; Uzun, L., Progress in conducting polymers for biointerfacing and biorecognition applications. Sensors and Actuators Reports 2021, 3, 100035.

31. Khodagholy, D.; Doublet, T.; Gurfinkel, M.; Quilichini, P.; Ismailova, E.; Leleux, P.; Herve, T.; Sanaur, S.; Bernard, C.; Malliaras, G. G., Highly conformable conducting polymer electrodes for in vivo recordings. Advanced Materials 2011, 23, (36), H268.

32. Ganji, M.; Kaestner, E.; Hermiz, J.; Rogers, N.; Tanaka, A.; Cleary, D.; Lee, S. H.; Snider, J.; Halgren, M.; Cosgrove, G. R., Development and translation of PEDOT: PSS microelectrodes for intraoperative monitoring. Advanced Functional Materials 2018, 28, (12), 1700232.

33. Pranti, A. S.; Schander, A.; Bödecker, A.; Lang, W., PEDOT: PSS coating on gold microelectrodes with excellent stability and high charge injection capacity for chronic neural interfaces. Sensors and Actuators B: Chemical 2018, 275, 382–393.

34. Venkatraman, S.; Hendricks, J.; King, Z. A.; Sereno, A. J.; Richardson-Burns, S.; Martin, D.; Carmena, J. M., In vitro and in vivo evaluation of PEDOT microelectrodes for neural stimulation and recording. IEEE transactions on neural systems and rehabilitation engineering 2011, 19, (3), 307–316.

35. Kozai, T. D.; Catt, K.; Du, Z.; Na, K.; Srivannavit, O.; Haque, R.-u. M.; Seymour, J.; Wise, K. D.; Yoon, E.; Cui, X. T., Chronic in vivo evaluation of PEDOT/CNT for stable neural recordings. IEEE transactions on biomedical engineering 2015, 63, (1), 111–119.

36. Khodagholy, D.; Gelinas, J. N.; Thesen, T.; Doyle, W.; Devinsky, O.; Malliaras, G. G.; Buzsáki, G., NeuroGrid: recording action potentials from the surface of the brain. Nature neuroscience 2015, 18, (2), 310–315.

37. Boehler, C.; Carli, S.; Fadiga, L.; Stieglitz, T.; Asplund, M., Tutorial: guidelines for standardized performance tests for electrodes intended for neural interfaces and bioelectronics. Nature protocols 2020, 15, (11), 3557–3578.

38. Vajrala, V. S.; Saunier, V.; Nowak, L. G.; Flahaut, E.; Bergaud, C.; Maziz, A., Nanofibrous PEDOT-Carbon Composite on Flexible Probes for Soft Neural Interfacing. Frontiers in bioengineering and biotechnology 2021, 9, 780197.

39. Castagnola, V.; Bayon, C.; Descamps, E.; Bergaud, C., Morphology and conductivity of PEDOT layers produced by different electrochemical routes. Synthetic metals 2014, 189, 7–16.

